# Overlapping Neural Representations for Dynamic Visual Imagery and Stationary Storage in Spatial Working Memory

**DOI:** 10.1101/2022.09.24.509255

**Authors:** Eren Günseli, Joshua J. Foster, David W. Sutterer, Lara Todorova, Edward K. Vogel, Edward Awh

## Abstract

Representations in working memory need to be flexibly transformed to adapt to our dynamic environment and variable task demands. Recent work has demonstrated that activity in the alpha frequency band enables precise decoding of visual information during both perception and sustained storage in working memory. Extant work, however, has focused exclusively on the representation of static visual images. Here we used EEG recordings to examine whether alpha-band activity supports the dynamic transformation of representations in spatial working memory using an imagery task that required the active shifting of a stored position to a new position. In line with recent findings, a common format of alpha-band activity precisely tracked both the initial position stored in working memory as well as the transformed position, with the latter emerging approximately 800-1200 ms following an auditory cue to “rotate” to a new position. Moreover, the time course of this transformation of alpha activity predicted between-subject differences in manual reaction time to indicate the new position (Experiment 1), as well as within-subject variations in saccade latency in a speeded version of the task (Experiment 2). Finally, cross-training analyses revealed robust generalization of alpha-band reconstruction of working memory contents before and after mental transformation. These findings demonstrate that alpha activity tracks dynamic transformations of representations in spatial working memory, and that the format of this activity is preserved across the initial and transformed memory representations. These findings highlight a new approach for measuring voluntary shifts in online memory representations and show common representational formats during dynamic mental imagery and the maintenance of static representations in working memory.

## Introduction

Working memory (WM) is the ability to store and manipulate information to guide behavior in ongoing tasks (Baddeley & Hitch, 1974). Most daily tasks require the effective functioning of WM. In line with this, successful WM functioning predicts higher-order cognitive skills such as reasoning, problem-solving, and intelligence (Carpenter et al., 1990; Chow & Conway, 2015; Engle et al., 1999; Fukuda et al., 2010; Kane & Engle, 2002). Since we live in a dynamic environment, the transformation of the contents of WM is essential for accurately guiding human behavior. For example, while driving, we regularly check the rear window mirror; between each glance at the mirror, we estimate the position of the car approaching from behind based on its position and speed the last time we checked the mirror. In other words, we transform its location in our minds. If the car in front of us stops abruptly and we have to make a rapid decision (e.g., whether to put the brakes hard or switch lanes) without the time to check the rear window, we can rely on this internally transformed spatial WM representation. Given the important role of WM transformation in daily life, exploring these processes is crucial not only for understanding the nature of WM but also for human behavior broadly.

Recently, activity in the alpha-band frequency has been shown to enable precise decoding of visual information during perception, sustained storage in WM (Foster et al., 2015, 2017), long-term memory retrieval (Sutterer et al., 2019), and stationary mental imagery (Xie et al., 2020). Moreover, cross-temporal analyses have shown that the same patterns of alpha-band activity generalize across perception and mental imagery suggesting that they rely on overlapping networks (Xie et al., 2020). However, these studies focused primarily on representations of static images. As in the driving example provided above, most daily-life functions require mentally shifting and transforming representations within WM. Thus, our goal here was to examine whether dynamic transformation of spatial representations during visual imagery is indexed by the same spatially selective alpha activity that tracks locations held in working memory. An alternative possibility is that internally directed transformations of spatial representations may alter the format of the representation. For instance, Yu et al. (2020) found evidence for strong transformations (i.e., “rotations”) of the format of neural representations when they were moved in and out of the focus of attention. Thus, we examined whether dynamic *transformation* of a representation in spatial working memory has similar effects as when an observer voluntarily switches attention from one item to another.To examine these questions, we used an inverted encoding model (IEM) in the EEG to reconstruct the contents of spatial WM in a time-resolved fashion during a mental transformation task that required mentally shifting a stored position to a new position. First, we tested whether the alpha-band activity would index dynamically shifting representations in spatial WM. Second, by training the IEM model trained on the EEG signal *before* transformation and testing it for the WM content *after* transformation, we examined whether there is a common format for transformed and static representations in visual WM.

We found that the contents of spatial WM could be reconstructed based on the topographic distribution of alpha-band power both for non-transformed and transformed WM. Moreover, the IEM model trained on the EEG signal before transformation successfully reconstructed the WM content after mental transformation, suggesting a common representational format for transformed and static WM representations as reflected in the alpha band topography. Finally, these patterns of alpha activity were sensitive to the efficiency with which subjects could indicate the transformed location, both between individuals (Experiment 1) and across trials within individuals (Experiment 2). These findings highlight a new approach for tracking the dynamic transformations of representations in visual WM in a time-resolved manner, and they show a clear correspondence between the representational formats that emerge during WM and mental imagery.

## Method

### Subjects

Fifty-six healthy volunteers participated in the experiments for monetary compensation ($15/hour), 37 in Experiment 1 and 19 in Experiment 2. The target number of subjects for Experiment 2 was determined based on a downsampling procedure using the data from Experiment 1 that tested how many data points were required to observe a significant CTF slope for the transformed location during the second retention interval (see Design and Procedure). Subjects reported normal or corrected-to-normal vision and provided informed consent according to procedures approved by the University of Chicago Institutional Review Board. Subjects were excluded from analysis if, after artifact rejection, they had less than 75 trials per angular location for either the initial position or the transformed position (seven in Experiment 1, zero in Experiment 2). One additional subject was excluded from the analysis in Experiment 2 because of an interrupted communication between the stimulus presentation computer and the EEG recording computer. In addition to the numbers provided above, in Experiment 2, 4 volunteers did not complete data collection (one battery died, two dropped out, one the eye tracker could not track eyes). Thus, the analysis included 30 subjects in Experiment 1 (age *Mean* = 23.2 *Standard deviation* = 4.1; 17 male) and 18 subjects in Experiment 2 (age *M* = 23, *SD* = 3.7; 7 male).

### Stimuli

Stimuli were generated in Matlab (Mathworks) using the Psychophysics Toolbox (Brainard 1997; Pelli 1997) and were presented on a 24-in. LCD monitor (refresh rate: 120 Hz). Viewing distance was ∼100 cm in Experiment 1 and ∼75 cm in Experiment 2. **Figure 1** shows an example trial. The background color was gray (22.5 Cd/m^2^). Throughout the trial, there was a black fixation circle (0.8 Cd/m^2^; radius ∼0.1° visual angle) at the center of the screen. Throughout the trial except for the sample display, five dark gray placeholders (9.8 Cd/m^2^; radius 0.8° in Experiment 1, radius 1.1° in Experiment 2) were presented centered equidistantly on an imaginary circle (radius 3° in Experiment 1, radius 5° in Experiment 2). During the sample display, one of the placeholders was replaced by a red circle (15.5 Cd/m^2^; radius 0.8° in Experiment 1, radius 1.1° in Experiment 2) that indicated the memory location. At the test display, there were light gray rings (43.9 Cd/m^2^; thickness ∼0.1°) around the placeholders. At the feedback screen, the outline of the placeholder that the subject selected was presented in white (124.7 Cd/m^2^) and the placeholder that contained the correct transformed position was filled in red as it was in the memory display. Additionally, during the feedback screen in Experiment 2 the accuracy of response was verbally indicated (i.e. “Correct”, “Wrong”, “You did not fixate on any placeholder”).

**Figure 1.**
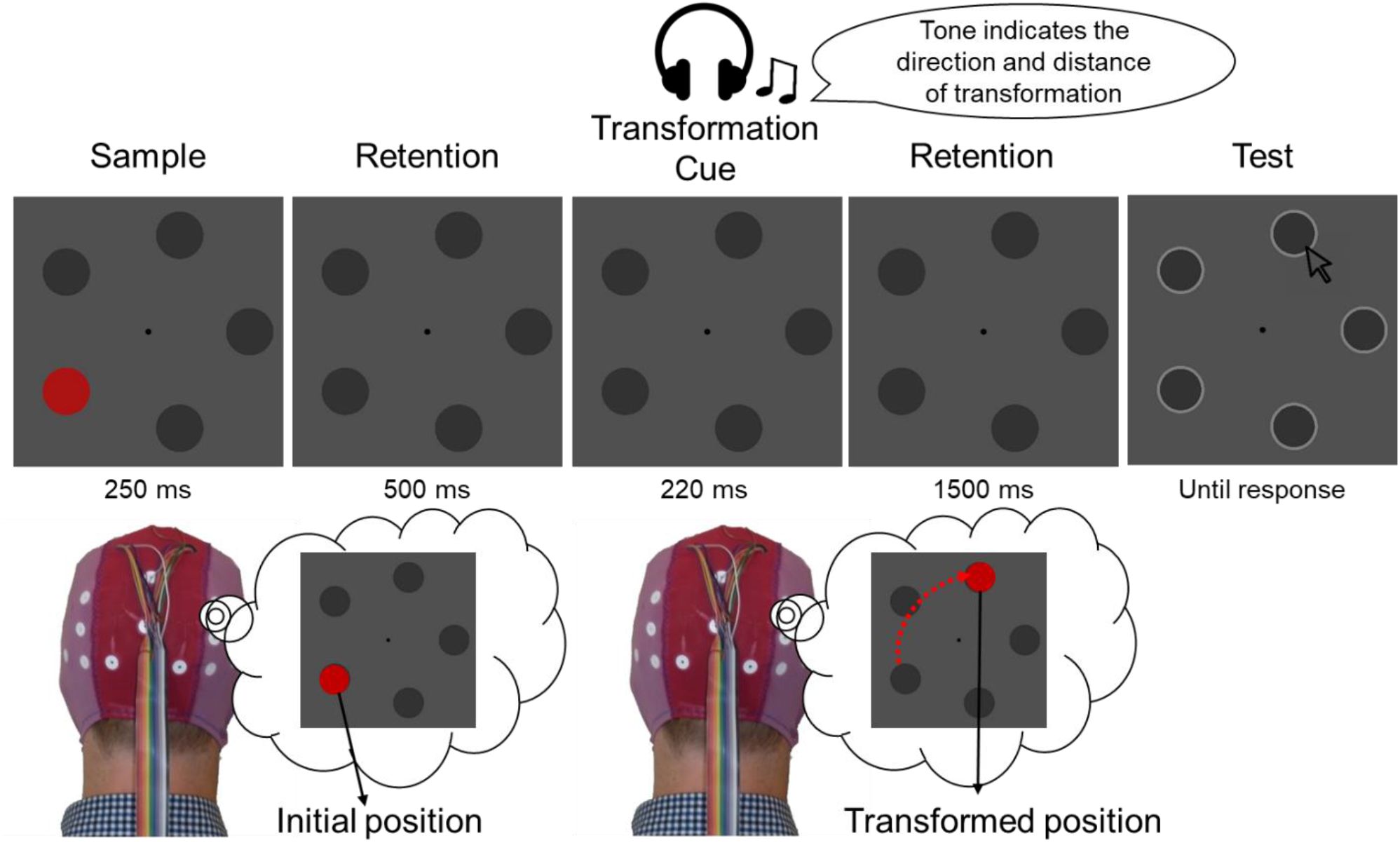
The experimental procedure in Experiment 1. Subjects were shown an initial position to remember. After a retention interval, they transformed the position in their mind based on an auditory transformation cue that indicated the direction and distance of the transformation via its pitch and repetition. For example, two beeps with a high pitch indicated a clockwise 2-step movement. At the end of the trial, subjects used mouse click to indicate which placeholder contained the transformed position. Experiment 2 was identical except the sample display was 150 ms, the second retention interval was 1000 ms, and the response was made by the gaze position recorded with an eye tracker instead of a mouse click.

The sound files for the transformation cue were created using Adobe Audition (http://www.adobe.com). The transformation cue was played through earphones (ER-3C audiometric earphone 10 ohms, www.etymotic.com). The tones were 174.61 Hz, 440 Hz, and 1108.73 Hz for the counterclockwise move, stay, and clockwise move, respectively. These frequencies correspond to notes in Western music that divide the octave into three equal steps. The single tones, which indicated a one-step movement, were normalized according to the equal-loudness contour. For two-step transformation cues, the middle of the tone was replaced with 80 ms of silence resulting in a double tone. All sound files were 220 ms. Single tones, which indicate a one-step transformation, have signal throughout the file, while double tones consist of two 70 ms signals interleaved with an 80 ms of silence in between. All sound files had 5 ms ramps at the beginning and end of the files to prevent artifacts during playback. In Experiment 2, each frequency was played in a different timbre (guitar for low, piano for mid, and flute for high frequency) to increase discriminability.

### Design and Procedure

Subjects initiated each trial by pressing the space bar. The fixation circle was presented for a randomly sampled duration between 600 and 1000 ms. Next, the memory display was presented for 250 ms in Experiment 1 and 150 ms in Experiment 2. Following the initial retention interval of 500 ms, the transformation cue was played through earphones for 220 ms during which the visual display was the same as the rest of the retention interval. After a second retention interval of 1500 ms in Experiment 1 and 1000 ms in Experiment 2, the test display was presented until response or an upper limit of 4000 ms. Following the response or 4000 ms, the feedback screen was presented for 1000 ms in Experiment 1 and 500 ms in Experiment 2. At the end of each block, written feedback was provided that indicated the block average accuracy in Experiment 1, and block average accuracy and RT in Experiment 2. In Experiment 2, the trial was aborted, and written feedback was presented for 500 ms if there was an ocular artifact during the trial. The ocular artifact feedback told the subject that they moved their eyes too early (i.e., before the test display) or blinked their eyes.

Subjects were instructed to remember the initial position until the transformation cue and to transform the position in their mind to the location indicated by the transformation cue as fast as possible following the cue. They were also told to hold the transformed position in mind until the test display. In Experiment 1, the task was to use the mouse to click on the placeholder that contained the transformed position as indicated by the transformation cue. In Experiment 2, subjects made the response by gaze position instead of a mouse click. Specifically, they were told to move their eyes to the placeholder that they thought contained the transformed position as fast as possible without risking accuracy. For both experiments, subjects were instructed to maintain fixation from the beginning of the trial until the onset of the test display.

At the beginning of each experiment, there was a cue familiarization phase of 50 trials at which subjects passively viewed the presentation of the initial position, and following the audio play of the transformation cue the presentation of the transformed position. This was to ensure that subjects learned the relationship between each tone and its meaning. After the cue familiarization, subjects performed a practice phase of 50 trials which were identical to the real experiment. The real experiment consisted of 16 blocks of 50 trials in Experiment 1 and 12 blocks of 125 trials in Experiment 2. Initial and transformed positions were counterbalanced using a full factorial design within each block. That is, each combination of the initial position and the transformed position was equally presented. After artifact rejection, the average number of trials used in analyses was 594.9 (*SD*= 91.9) in Experiment 1 and 1304.2 (*SD*= 147.5) in Experiment 2.

### EEG Recording

We recorded EEG activity using 30 active Ag/AgCl electrodes mounted in an elastic cap (Brain Products actiCHamp, Munich, Germany). We recorded from International 10-20 sites: FP1, FP2, F7, F3, Fz, F4, F8, FC5, FC1, FC2, FC6, C3, Cz, C4, CP5, CP1, CP2, CP6, P7, P3, Pz, P4, P8, PO7, PO3, PO4, PO8, O1, Oz, and O2. Two additional electrodes were placed on the left and right mastoids, and a ground electrode was placed at position FPz. All sites were recorded with the left-mastoid reference and were re-referenced offline to the algebraic average of the left and right mastoids. Electrophysiological signals were filtered (low cut-off = .01 Hz, high cut-off = 80 Hz, slope from low-to high-cutoff = 12 dB/octave) and recorded with a 500 Hz sampling rate. Impedance values were kept below 15 k Ω. Eye movements and blinks were monitored using EOG, recorded with passive Ag/AgCl electrodes. Horizontal EOG was recorded from a bipolar pair of electrodes placed ∼1 cm lateral to the external canthi. Vertical EOG was recorded from a bipolar pair of electrodes placed ∼2 cm above and below the right eye.

## Data Analysis

### Behavioral

Only trials with accurate responses were used for reaction time (RT) analysis. For the investigation of the relationship between behavioral and EEG measures, only artifact-free EEG trials were used to match the trials that are used to calculate behavioral and EEG scores.

### Eye-tracking

Gaze position was monitored using a desk-mounted infrared eye-tracking system (EyeLink 1000 Plus, SR Research, Ontario, Canada), operating in the remote mode in Experiment 1 and the chin-rest mode in Experiment 2. According to the manufacturer, this system provides a spatial resolution of 0.01° and an average accuracy of 0.25-0.5°. The gaze position was sampled at 500 Hz. We obtained usable eye-tracking data for 26 out of 30 subjects in Experiment 1. In Experiment 2, the behavioral response was made by ocular position. Therefore, the experiment was aborted for one subject whose eye tracker data was not stable. All the remaining subjects had usable eye-tracking data. In Experiment 2, online feedback regarding ocular artifacts was provided. Trials were aborted if the eyes deviated away from the fixation by 1.6° visual degrees from the sample display until the test display. This threshold was adjusted during data collection based on the eye tracker data quality to optimize ocular artifact detection. For unstable gaze data, the threshold was increased to detect as many real eye movements as possible and to minimize unnecessary trial abortions due to eye tracker noise. Whereas, for stable gaze data the threshold was decreased to detect as many eye movements as possible. In Experiment 2, the responses were registered via gaze position recording during the test display. At the test display, subjects moved their eyes to the placeholder that they thought contained the transformed position. A response was registered if the average gaze position of both eyes were within an imaginary radius of 1.6° visual degrees around the center of any placeholder. This threshold, similar to the ocular artifact threshold, was adjusted during data collection based on the quality of the eye tracker data to maximize accurate response registration.

## Artifact Rejection

The artifact rejection was performed by visual inspection of the EEG signal for recording artifacts (blocking, muscle noise, and skin potentials), and the EOG signal and eye-tracking data (for those subjects with usable eye-tracking data) for ocular artifacts (blinks and eye movements). Trials with an inaccurate response were excluded from all EEG analysis except the investigation of the relationship between EEG and response accuracy.

### Power Analyses

Power analyses were performed using Matlab’s Signal Processing toolbox (Mathworks, Natick, MA), EEGLAB toolbox (Delorme & Makeig, 2004), and FieldTrip toolbox (Oostenveld et al., 2011). To isolate frequency-specific activity at each electrode, the raw EEG signal was bandpass filtered using a Butterworth filter as implemented by the ft_preproc_bandpassfilter.m function of FieldTrip Toolbox (Oostenveld et al., 2011). A Hilbert Transform (Matlab Signal Processing Toolbox) was applied to the bandpass-filtered data, to extract a complex analytic signal, 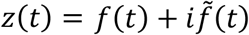, of the band-pass filtered EEG, *f*(*t*), where 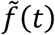 is the Hilbert Transform of *f*(*t*) and 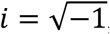, using the following Matlab syntax:

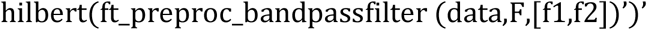

where data is the raw EEG (trials x samples), F is the sampling frequency (500 Hz), f1 and f2 are the boundaries of the frequency band to be isolated. For the time-frequency analysis, we f1 and f2 spanned across 4 Hz to 50 Hz in increments of 2 Hz with a 2-Hz bandpass: f1 and f2 were 4 and 6 to isolate 4-to 6-Hz activity; 6 and 8 to isolate 6-to 8-Hz activity; and so on. For alpha-band analyses, f1 and f2 were 8 and 12 Hz, respectively.

### Inverted Encoding Model of Spatial Position

#### Reconstructing the content of spatial WM

Following Foster et al. (2016), we used an IEM to reconstruct spatially selective CTFs from the scalp distribution of total power across electrodes. We assumed that power measured at each electrode reflects the weighted sum of five spatial channels (i.e., neuronal populations), each tuned for a different angular location (c.f. Foster et al., 2016). We modeled the response profile of each spatial channel across angular location bins as a half sinusoid raised to the seventh power, given by:

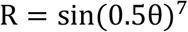

where *θ* is the angular location (ranging from 0° to 359°), and *R* is the response of the spatial channel in arbitrary units. This response profile was circularly shifted for each channel such that the peak response of each spatial channel was centered over one of the five positions corresponding to the placeholder positions (0°, 72°, 144°, 216°, 288°, see **Figure 1**).

An IEM routine was applied to each time point. This routine proceeded in two stages (train and test). In the training stage, training data *B*_*1*_ were used to estimate weights that approximate the relative contribution of the five spatial channels to the observed response measured at each electrode. Let *B*_*1*_ (*m* electrodes × *n*_*1*_ observations) be the power at each electrode for each measurement in the training set, *C*_*1*_ (*k* channels × *n*_*1*_ observations) be the predicted response of each spatial channel (determined by the basis functions) for each measurement, and W (*m* electrodes × *k* channels) be a weight matrix that characterizes a linear mapping from “channel space” to “electrode space”. The relationship between *B*_*1*_, *C*_*1*_, and *W* can be described by a general linear model of the form:

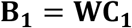

The weight matrix was obtained via least-squares estimation as follows:

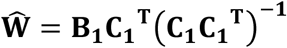

In the test stage, with the weights in hand, we inverted the model to transform the observed test data *B*_*2*_ (*m* electrodes × *n*_*2*_ observations) into estimated channel responses, *C*_*2*_ (*k* channels × *n*_*2*_ observations):

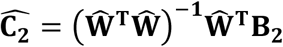

Each estimated channel response function was circularly shifted to a common center (i.e., 0° on the “Channel Offset” axes of the plots in **Figure 2**) by aligning the estimated channel responses to the channel tuned for the stimulus location to yield CTFs. The IEM routine was performed separately for each time point.

**Figure 2.**
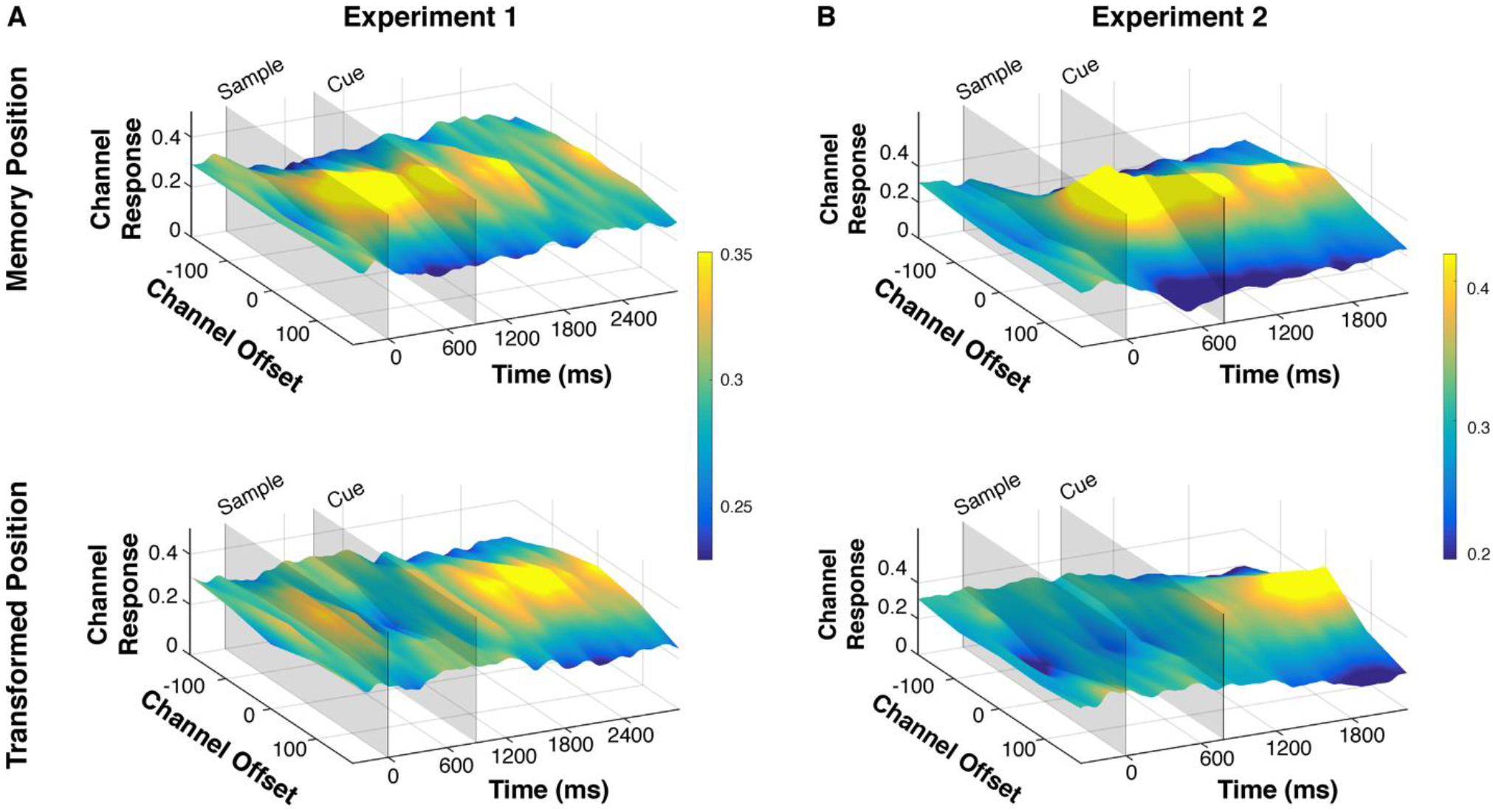
Time-resolved channel tuning functions (CTFs) reconstructed using an IEM on the alpha-band (8-12 Hz) power in the EEG signal tracked initial and transformed positions both in **(a)** Experiment 1 and **(b)** Experiment 2. CTFs are shown separately for the initial position (top row) and transformed position (bottom row). The onset of sample and cue displays are shown with gray rectangles. Higher and yellower peaks in the CTFs indicate larger location selectivity in alpha-band power. In both experiments, alpha-band power showed location selectivity for the initial position after the sample display and for the transformed position after the transformation cue.

We used a “leave-one-out” cross-validation routine such that two blocks of estimated power values (see Alpha-Band Power Analysis) served as *B*_*1*_ and were used to estimate, *Ŵ* and the remaining block served as *B*_*2*_ and was used to estimate *C*_*2*_, ensuring that the training and test data were always independent. This process was repeated until each of the three blocks were held out as the test set, and the resulting CTFs were averaged across each test block. This IEM routine was applied separately for each subject, and statistical analyses were performed on the reconstructed CTFs.

To prevent bias in our analysis, we equated the number of observations across position bins within each block. Specifically, we calculated the minimum number of trials for any given position bin *n* for each subject and assigned *n*/3 many trials for each position bin to each of the three blocks. Importantly, the blocks were independent (i.e., no trial was repeated across blocks) to prevent circularity in the cross-validation procedures used for the IEM routine (see Forward Encoding Model). Total power was then calculated for each position bin for each block, resulting in a *p*\**b* × *m* × *s* matrix, where *p* is the number of position bins, *b* is the number of blocks, *m* is the number of electrodes, and *s* is the number of time samples.

Finally, because we equated the number of trials across position bins within blocks, a random subset of trials was not included in any block. We randomly generated 30 block assignments each resulting in a *p*b × m × s* power matrix. The IEM routine was applied to the matrices of power values for each block assignment, and their outputs (i.e., channel response profiles) were averaged. This iterative approach better utilized the complete data set for each subject and reduces noise in the resulting CTFs by minimizing the influence of idiosyncrasies in estimates of power specific to any given assignment of trials to blocks.

#### Testing the representational similarity between the WM content for transformed and pre-transformed positions

The IEM analysis described above enabled testing whether there is location-selective information in the EEG signal for a given time point, separately for the initial position and the transformed position. To test whether the storage of the pre-transformed content and the transformed content share an overlapping representational space, we performed the aforementioned IEM with the following difference: We trained an IEM for the initial position using data averaged across a pre-cue time and tested the IEM for the transformed position for each time point. To determine the time to be used for training the IEM, first, we ran a cross-temporal IEM for the initial position until the cue onset. The CTF slope for this IEM was then visually inspected to assess how long it took for the CTF, thus the WM representation, to stabilize. Based on visual inspection, we decided that the CTF slope was stabilized after ∼300ms for both experiments (see **Supplementary Figure 1**). In order to minimize the contamination in the EEG signal caused by mental transformation, we set the end of the training time window to be 50 ms before the onset of the transformation cue. Thus, we trained the IEM on the initial position using the alpha-band power averaged from 300 ms till 700 ms in Experiment 1, and from 300 ms till 600 ms in Experiment 2. Then, we tested the IEM on the transformed position at each time point. Thus, the model was blind to the transformed location information since it was trained on the initial position using data before the transformation cue. Therefore, we hypothesized that this analysis would reveal location-selective information for the transformed position only if the storage of the *transformed* WM content relies on an alpha-band power topography that is similar to the storage of the *initial*, pre-transformed WM content.

### Statistical Analysis

To quantify the location selectivity of the CTFs we estimated the CTF slope (i.e., the “peakedness” of the channel response at channel offset zero relative to farther channel offsets). CTF slope was estimated using linear regression by collapsing the channel responses across channels that were equidistant. To test whether the CTF slope was statistically above chance, we tested whether the CTF slope was different from zero using a one-sample t-test (two-tailed).

Because the mean CTF slope may not be normally distributed under the null hypothesis, we employed a Monte Carlo randomization procedure to empirically approximate the null distribution of the t-statistic.

Specifically, we implemented an IEM by randomizing the location labels within each block so that the labels were random with respect to the observed EEG signal in each electrode. This randomization procedure was repeated 1,000 times to obtain a null distribution. CTF slope at each time point was compared against this null distribution using a two-tailed t-test. Multiple comparisons correction was established using cluster-based permutation testing. First, a difference score was calculated by subtracting the surrogate null distribution from the real CTF slopes and this difference score was tested against zero using a two-sample t-test. Four or more temporally adjacent data points with a p-value smaller than 0.05 were clustered together. Then, signs of the difference scores were randomly shuffled across participants 1,000 iterations. At each iteration, t-tests were performed on shuffled data. A cluster-level statistic was calculated by taking the sum of the t-values within each cluster, separately for positive and negative clusters. The p-value for each cluster was calculated as the number of times the sum of the absolute t-values within the cluster under random shuffling exceeds that of the t-values within the observed cluster. A cluster was considered significant if the calculated p-value was smaller than 0.05. This approach corrects for multiple comparisons by taking into account clusters of modulations in data that can be expected by chance (Maris & Oostenveld, 2007). The same approach was used for the statistical analyses of IEM across multiple frequency bands, ranging from 4 Hz to 50 Hz (4-6 Hz, 6-8 Hz, … 48-50 Hz), except that this time we downsampled the data to 50 Hz (i.e., one sample every 20 ms) prior to statistical analyses to reduce the computation time.

### EEG-Behavior Relationship

To test whether the location selectivity of the CTF for the transformed WM content predicts behavior, in Experiment 1 we looked at the correlation between the average CTF slope for the transformed content following the transformation cue and the grand average accuracy and RT across subjects. For this analysis, we matched the trials used to calculate behavioral measures to the trials used to calculate CTF slopes by including only artifact-free trials. Also, we equated the number of trials across subjects to match the subject with the lowest number of trials to eliminate variance in mean CTF slope across subjects that is due to differences in EEG noise. Specifically, the number of trials in each position bin in each block was set to be 86 for the correlation with accuracy, and 76 for the correlation with RT, which was the minimum across subjects for all trials and correct response trials, respectively. Moreover, to capture the variability in RT for the transformation of WM contents, we only included ‘move’ trials (i.e., excluded ‘stay’ trials) in this analysis. In Experiment 2, the variability in RT was very low, but the number of artifact-free trials was more than twice that in Experiment 1. Thus, we investigated the CTF-behavior relationship across trials instead of across subjects. We applied a median-split on mean RT per subject and performed the standard IEM approach described above separately for the above-median RT and below-median RT trials. Like in Experiment 1, only correct-response and move trials were included in this analysis.

### Eye Bias Control Analysis

To test whether changes in EEG voltage that are due to eye movements account for the location selectivity in the EEG signal, we ran the main CTF analysis described above with the following differences. We performed this analysis for Experiment 2, in which there was reliable eye-tracking data for every participant and trial. The position bins were defined based on eye positions instead of the actual initial and transformed positions. The initial position was defined as the location of the placeholder closest to the eye position averaged from the onset of the memory display until the onset of the transformation cue (i.e. 0-650 ms). The transformed position was defined as the location of the placeholder closest to the eye position averaged from the offset of the transformation cue until the onset of the probe (i.e. 870-1870 ms). Moreover, trials with noisy eye-tracking data were removed from this analysis resulting in fewer trials used compared to the main analysis, which resulted in the removal of only 2.44 trials on average (minimum 0, maximum 14 trials per subject). One subject who had less than 30 trials for a position bin in a ‘block’ of the CTF analysis was excluded from the analysis. Otherwise, the analysis was identical to our main analysis described above in Reconstructing the Content of Spatial WM. To ensure the validity of the comparison of the location selectivity across CTFs obtained based on WM location bins and eye position location bins, we performed the standard CTF analysis using the same subjects and trials used in the eye bias control analysis. If eye movements were responsible for the reconstruction of the transformed position our main analysis, then using position labels based on eye positions should result in location selectivity.

To statistically confirm that eye bias is not responsible for the CTF location selectivity for initial and transformed positions, we compared the CTF slopes for initial vs. transformed positions separately for the CTF analyses that are based on actual positions and eye positions. We hypothesized that if eye movements do not underlie the location selectivity in the EEG signal then a CTF slope difference between initial and transformed positions should be observed for actual position bins, but not eye-position-based bins.

## Results

### Behavior

Behavioral results show that subjects were able to perform the task with a high accuracy in both experiments. The grand average accuracy was 97.1% (*SD* = 2.6%) in Experiment 1 and 93.9% (*SD* = 4.9%) in Experiment 2. The response times were smaller and less variable in Experiment 2 (see Discussion). On average, the RT was 911 ms (*SD* = 363.7 ms) in Experiment 1 and 356.4 ms (*SD* = 73.7 ms) in Experiment 2.

### EEG

#### Alpha-band scalp topography tracks transformed content in spatial WM

**Figure 2**. shows the CTFs, and **Figure 3** shows the slopes of the CTFs for both experiments. Our analysis showed that the initial memory position was actively maintained throughout the initial retention interval until the transformation cue for both experiments, as alpha-band power CTFs revealed a sustained spatial selectivity during this interval. This finding replicates Foster et al. (2016) and shows that alpha-band scalp topography tracks the contents of spatial WM.

Importantly, the location selectivity for the *transformed* position was also tracked in the alpha-band topography. It emerged following the transformation cue and was sustained until the test display. This result extends previous findings and shows that alpha-band topography can be used to reconstruct *transformed* contents of spatial WM in addition to the non-transformed contents. Foster et al. (2016) found that reconstruction of the contents of spatial WM is specific to the alpha-band and is not reflected in other frequency bands. Next, we tested if the same range of frequency bands tracks the transformed contents of WM.

**Figure 3.**
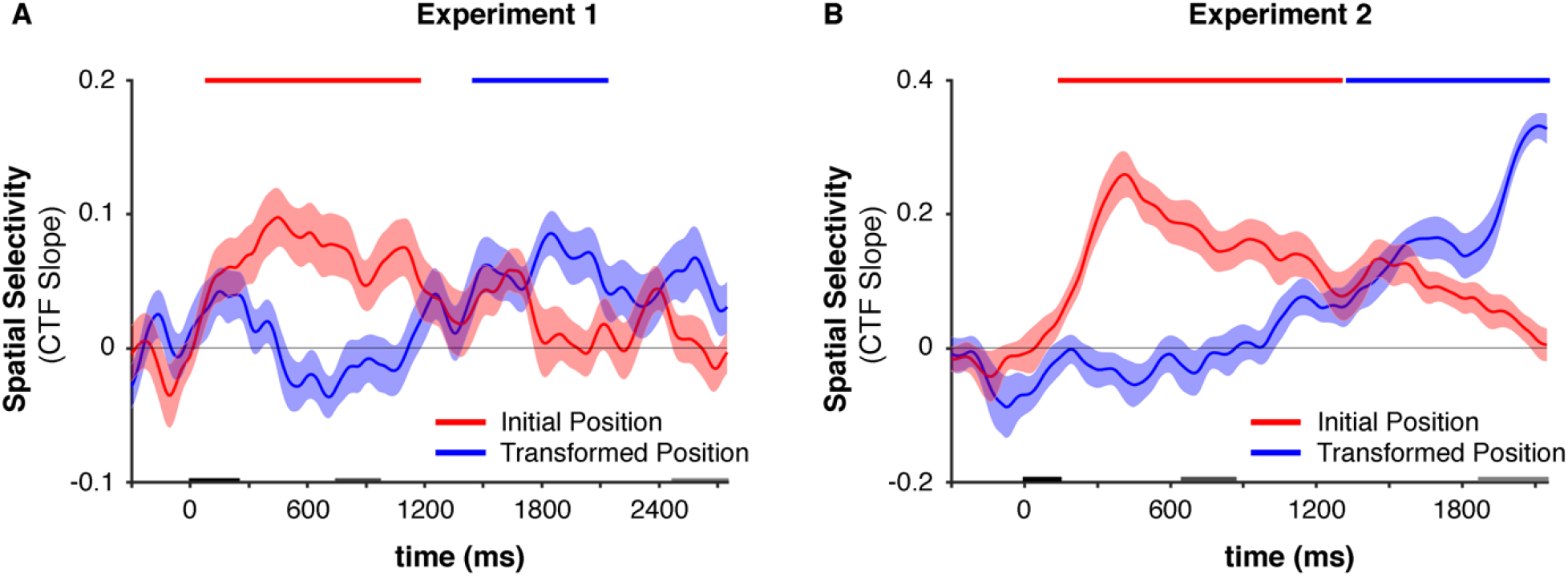
Location selectivity in the EEG that is quantified as CTF slopes for **(a)** Experiment 1 and **(b)** Experiment 2 shown in different colors for the initial and transformed positions. The markers along the top of the panes, shown in the same colors as the main CTF slopes, indicate the time points at which the CTF slope was reliably above chance as indicated by a cluster-based permutation test against a randomly permuted sample (p*<0*.*05;* two-tailed). Shaded error bars show the bootstrapped standard error of the mean.

#### Manipulated and pre-transformed content of WM are both represented in the alpha-band activity

Previous work has shown that WM representations can be represented in different frequency bands depending on their attentional status within WM (Rose et al., 2016). Thus, we examined whether transformed representations in WM were represented within the same range of frequency bands as the positions cued by external stimuli. To test this, we calculated CTFs across frequency bands ranging from 4 Hz to 50 Hz. For both experiments, location selectivity was mainly restricted to the alpha-band, both for the initial and transformed position (**Figure 4**). There are three exceptions to this. First, early in the trial, location selectivity was also present in theta band activity (∼ 4-7 Hz). This result is consistent with previous work that shows early (0-500 ms) location selectivity in the theta band for spatial WM representations (e.g., Foster et al., 2016). Importantly, this low-frequency location selectivity was not sustained either in the present work or in Foster et al. (2017), which suggests that it likely represents the initial encoding of stimuli instead of storage in WM. In line with this, there was no theta-band selectivity for the transformed position. Second, only in Experiment 1, there was a significant cluster approximately between 1500 to 2300 ms in a high-frequency band (∼40-50 Hz). However, this cluster had a negative CTF slope indicating a location ‘unselectivity’ for the initial position. This result is likely a false positive, because the same pattern was not observed in Experiment 2 which was much better powered because a larger number of trials were run. Third, in Experiment 2, right before the probe onset, there was location selectivity across all frequencies. Given that in Experiment 2, participants indicated their response by moving their eyes to the placeholder of the transformed position, we suggest that this broad-band location selectivity specific to the transformed position reflects eye-movement preparation related activity and not WM storage. In line with this, such frequency unspecific reconstruction was not observed in Experiment 1. Moreover, only the alpha-band location selectivity was common to both experiments, reaching significance approximately 300 ms after the offset of the transformation cue, which is approximately 550 ms before the broadband selectivity. Thus, we conclude that spatial WM representations were restricted to the alpha-band activity both for stationary and transformed contents.

**Figure 4.**
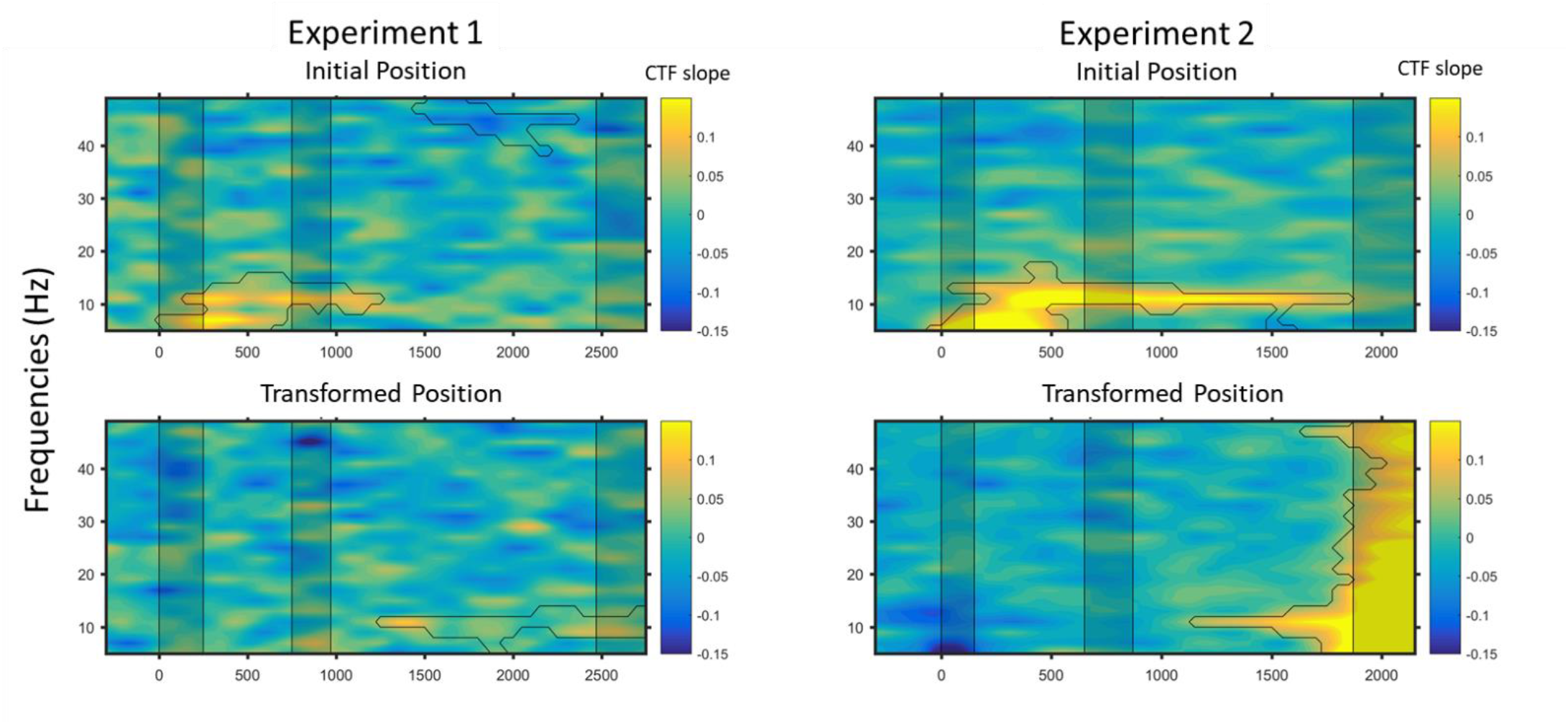
Location selectivity (i.e., CTF slope) in the EEG across frequency bands ranging from 4 Hz to 50 Hz quantified as CTF slopes for Experiment 1 (left) and Experiment 2 (right) shown in separate panels for initial (top) and transformed positions (bottom). The black lines indicate the time-frequency bands at which the CTF slope was reliably above chance as indicated by a cluster-based permutation test against a randomly permuted sample (p<0.05; two-tailed). Shaded rectangles show the duration within which the sample, transformation cue, and the probe were presented, respectively.

#### Transformed and pre-transformed content of WM share an overlapping representational space

Our results above show that alpha-band topography tracks both the initial (i.e., non-transformed) and transformed contents of spatial WM. A remaining question is whether the storage of transformed contents relies on overlapping neural mechanisms with the storage of non-transformed contents. Even though our main analysis mentioned above showed that transformed and non-transformed contents can be reconstructed based on alpha-band power topography, it is possible that the topographic distributions for transformed and non-transformed WM contents are different. This is because each CTF was constructed using separate IEMs that used different locations. The CTF for the initial position was reconstructed using an IEM that used the initial position for training and testing. Likewise, the CTF for the transformed position was reconstructed using an IEM that used the transformed position. To test whether the storage of transformed content in WM shares an overlapping representational space with the storage of pre-transformed content in WM, we trained the IEM for the initial position during the initial retention interval and tested the IEM for the transformed position at each time point. Thus, the training of the IEM was blind to the information regarding the position of the transformed content. We hypothesized that this cross-training should demonstrate location selectivity only if the storage of initial and transformed contents in spatial WM rely on overlapping neural mechanisms as reflected in alpha-band power scalp topography. On the other hand, if the storage of transformed contents relies on a different pattern of neural activity than the initial storage in WM, then training the IEM on the initial position should not reveal location selectivity for the transformed position.

For both experiments, training the IEM based on the EEG activity pattern for the initial WM content prior to the transformation cue and testing it for the transformed content at each time point produced an accurate and reliable reconstruction of the transformed content following the transformation cue (**Figure 5**). This result suggests that the storage of the initial (i.e., pre-transformed) content and the transformed content in spatial WM share an overlapping neural representational space.

**Figure 5.**
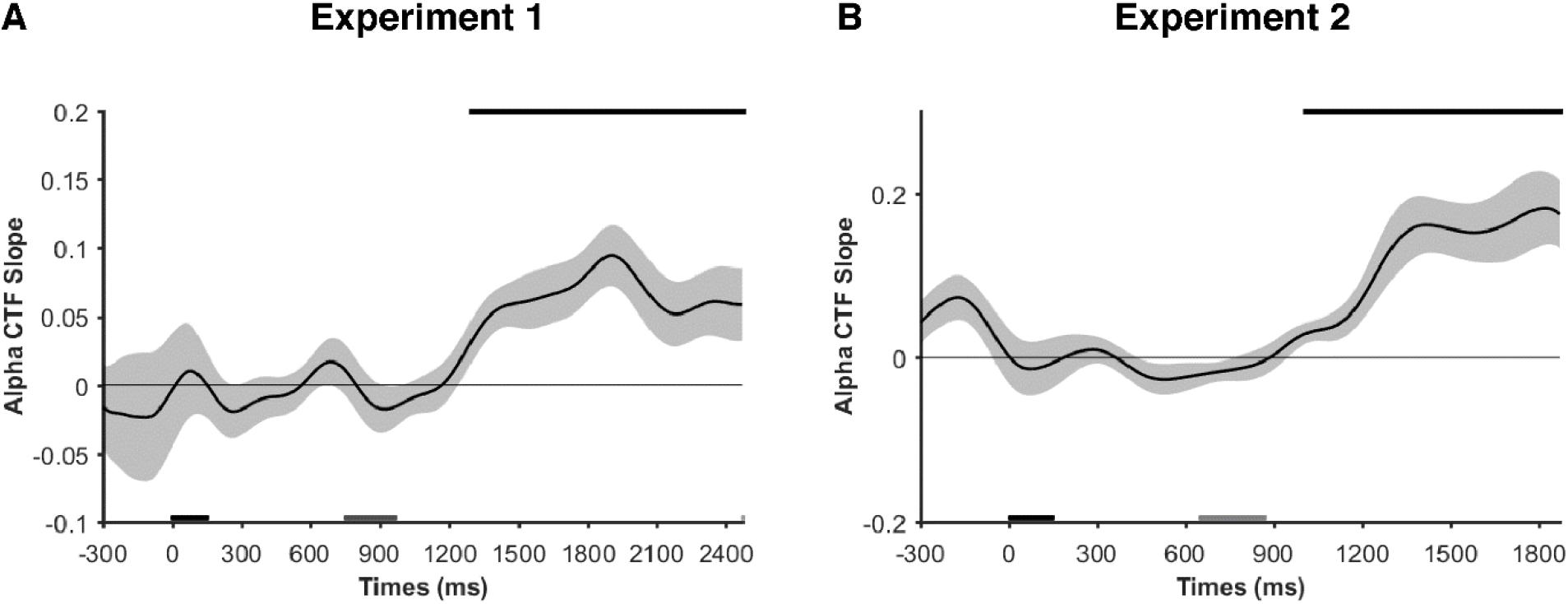
CTF slope for the *transformed* position obtained by cross-training the IEM based on the *initial* position *prior* to the cue and testing it for the transformed position at each time point. Shaded error bars show the bootstrapped standard error of the mean. The black markers along the top of the panes indicate the time points at which the CTF slope was reliably above chance as indicated by a cluster-based permutation test against a randomly permuted sample (p<0.05; two-tailed). Thus, transformed and non-transformed spatial WM contents have overlapping neural representations.

#### There is no evidence for analog mental transformation

On the two-step move trials, the mental transformation involved a placeholder between the initial and transformed positions. This allowed us to test whether the transformation was analog or discrete. As in the above section, we trained the IEM for the initial position during the initial retention interval. However, this time we tested it for the position between the initial and transformed positions at each time point. For this analysis, we only used two-step move trials, as one-step move and no-move trials did not involve an in-between position. If participants continuously shifted the mental image from the initial position to the transformed position, we should observe location selectivity for the placeholder in between the two. On the other hand, if transformation takes place in a discrete manner, then we should observe the waxing of the transformed position and the waning of the initial position in the absence of any location selectivity in the middle position. Our results support the discrete mental transformation model, as the location selectivity did not differ from zero at any time point during the retention interval following the transformation cue (**Figure 6**).

**Figure 6.**
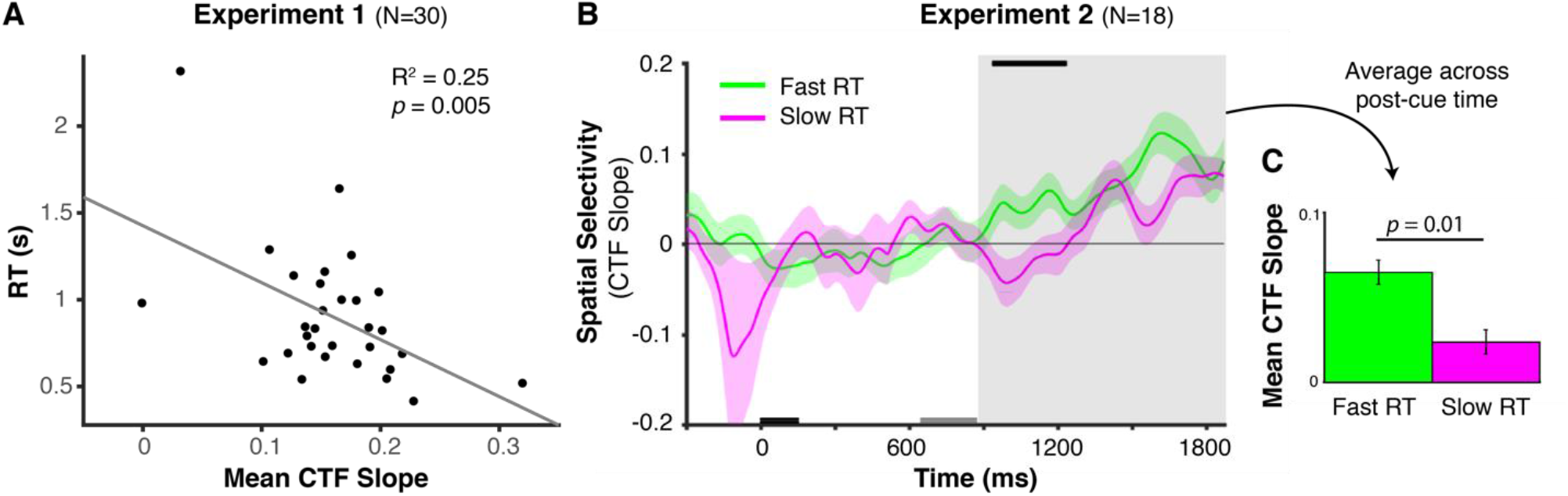
Higher location selectivity in the EEG predicted faster responses. **(a)** In Experiment 1, the mean CTF slope for the transformed position following the transformation cue predicted average RT across subjects (*p* = 0.005). **(b)** In Experiment 2, the CTF slope for the transformed position across time points given separately for fast response (i.e. below-median RT) and slow response (above median RT) trials shown in different colors. The black markers along the top of the pane indicate the time points at which the CTF slope was reliably different in fast response and slow response trials as indicated by a cluster-based permutation test (*p*<0.05; two-tailed). The dark gray and light gray bars on the x-axis indicate the timing of the sample and cue displays respectively. The big shaded gray region shows the 2^nd^ retention interval, from the offset of the transformation cue until the onset of the test display, at which the average CTF slope for the transformed position was calculated. Shaded error bars show the bootstrapped standard error of the mean. **(c)** The average CTF slope during the 2^nd^ retention interval (shown in the gray region in Figure 6b) separately for fast RT and slow RT trials. Fast RT trials had a larger CTF slope than slow RT trials (*p* = 0.01). The error bars show the standard error of the mean for data normalized for the between-subjects variance.

#### Alpha-band scalp topography for the transformed content predicts behavior

To examine whether alpha band activity was linked with behavior, we tested whether the selectivity of the CTF tracking the transformed representation was corelated with the reaction time for reporting the transformed position. Indeed, the average CTF slope for the transformed position after the transformation cue (i.e., from the offset of the transformation cue until the onset of the test display) predicted the average RT across subjects, R^2^ = 0.25, *p* = 0.005 (**Figure 6a**). There was no CTF slope – accuracy correlation, presumably because of the low variance in accuracy data (*M* = 97.1%. *SD* = 2.6%). Interestingly, this correlation was observed in Experiment 1 even though subjects were not told to prioritize speed when they made their manual responses. In Experiment 2, by contrast, subjects were instructed to move their eyes to the transformed position as quickly as they could after the onset of the test display, yielding much shorter response latencies. In addition, we collected twice the number of trials in Experiment 2 compared to Experiment 1, providing adequate data to look at CTF selectivity across fast and slow trials in a within-subject analysis. Thus, the data were divided based on a median split of saccade latencies. First, separate IEMs were used to calculate separate CTFs for above-median RT (i.e., slow response) and below-median RT (i.e. fast response) trials. As in Experiment 1, we averaged the CTF slope for the transformed position following the transformation cue. The average CTF slope for the transformed position was larger in fast response trials (*Mean Response Time* = 202.3 ms; *SD* = 26.0 ms), *M* = 0.06 (*SD* = 0.06), compared to slow response trials (*Mean Response Time* = 510.3 ms; *SD* = 130.5 ms), *M* = 0.02 (*SD* = 0.05), *t*(17), 2.86, *p*=0.01 (**Figures 6b and 6c**). Together, these results suggest that the active storage of the transformed content in WM, as indexed by the larger CTF slope, predicted faster eye movements to the transformed position.

#### Eye movements do not account for the location selectivity in the EEG signal

Trials with ocular artifacts were removed prior to any analysis. However, it is possible that subtle but systematic eye movements resulted in changes in scalp topography that underlies our CTF location selectivity results. To test this alternative explanation, we repeated our main analysis using recorded eye positions to determine the position bins in the IEM. We performed this analysis for Experiment 2 for which we had reliable eye-tracking data for all participants and almost every trial, as trials with eye movements were aborted and repeated in Experiment 2. This analysis did not reveal a clear location selectivity for either the initial position nor the transformed position in WM (**Supplementary Figure 2**). Moreover, the location selectivity shift from the initial position before the cue to the transformed position after the cue was observed only for the CTF slopes calculated based on the IEM that used actual position bins, but not the position bins at which the gaze position was closest to. Thus, eye movements cannot explain our EEG results.

## Discussion

The transformation of WM representations is crucial for humans to successfully guide their behavior in a dynamic world. Here, we extended prior work showing that stationary spatial WM and mental imagery representations can be reconstructed based on the topography of alpha-band activity (8-12 Hz) (Foster et al., 2015, 2017; Xie et al., 2020). Specifically, we show that alpha-band activity also indexes dynamic transformations of representations in spatial WM, with spatially-selective activity tracking the transformed positions within several hundred milliseconds of the transformation cue. Successful reconstructions were specific to the alpha-band power for both transformed and initial representations. In addition, cross-training analyses showed that the format of alpha activity was similar across the transformed and initial positions. Specifically, training the IEM based on the *initial* WM content before the transformation cue was sufficient to track the *transformed* WM position from the subsequent phase of the trial. Finally, the location selectivity for the transformed position in the EEG signal predicted which subjects reported that location the fastest (Experiment 1) and distinguished between fast and slow responses in a within-subject analysis (Experiment 2). Thus, these findings highlight a new approach for precisely tracking dynamic transformations of WM representations, and reveal a common format of alpha activity for static and dynamic representations.

Recent evidence suggests that alpha-band activity in the EEG enables the tracking of visual information during stationary storage in WM (Foster et al., 2015, 2017) and mental imagery (Xie et al., 2020). In a mental imagery task, Xie et al. (2020) provided an auditory recording of a word that represents an object (e.g., apple) that participants were instructed to visually imagine. A classifier was trained on the data when participants viewed the same objects on the screen and tested when participants were given auditory cues to imagine the objects. EEG activity in the alpha-band power provided successful decoding of object categories suggesting that alpha-band activity underlies the perception and mental imagination of information in visual WM. However, the mental imagery task always followed the perception task where individuals viewed the same images across dozens of trials. Thus, it is likely that individuals retrieved these images from long-term memory when instructed to imagine them, as opposed to internally generating mental imagery. Moreover, the cross-decoding in Xie et al. (2020) was between perception and imagery, while we obtained it between initial static storage in WM and later dynamically transformed representation via mental imagery. Lastly, previous studies were focused on static representations in WM, either based on a visual presentation (Foster et al., 2015, 2017) or auditory retrieval cues (Xie et al., 2020). Thus, a key extension offered by the present work is the demonstration of overlap in the format of alpha activity during WM and imagery that is maintained during the dynamic transformations of spatial representations that were required by a mental transformation task.

Transformation of information has been previously studied using fMRI in the domain of mental imagery. These studies showed that motor areas, superior and inferior parietal cortex, and early visual areas are involved in mental rotation and transformation of information in VWM (Jordan et al., 2001; Kaas et al., 2010; Klein et al., 2000; Knauff et al., 2000; Koenigs et al., 2009; Richter et al., 2000). More recently, MVPA was used to investigate the brain regions that represent the contents of mental rotation. Subjects were given an orientation or a shape to remember and then they rotated the orientation in their mind based on rotation cues that indicated direction and degrees of rotation (e.g., clockwise 60°). The results showed that early visual areas (V1-V4) stored the initial orientation and also, following the rotation cue, the rotated orientation (Albers et al., 2013; Christophel et al., 2015). Moreover, Albers et al. (2013) obtained cross-decoding between training on WM retention and testing on mental imagery. Thus, these studies provide evidence for a neural overlap for storing stationary and mentally manipulated objects in WM. However, given the differences in cognitive and neural mechanisms for representing object vs. spatial information in WM both during storage and transformation (Carlesimo et al., 2001; Farah et al., 1988; Foster et al., 2017; Hecker & Mapperson, 1997; Logie & Marchetti, 1991; Luzzatti et al., 1998; Mecklinger & Pfeifer, 1996; Mohr & Linden, 2005; Smith et al., 1996, p. 19, 1996; Vecchi & Cornoldi, 1999; Vicari et al., 2006), it remained an open question whether transformation and storage of *spatial* information in WM also rely on overlapping neural representations. In addition, oscillatory activity measured with EEG may tap into distinct aspects of WM maintenance than does BOLD activity measured with functional MRI. Thus, our demonstration that oscillatory activity indexes the storage of both static and dynamic representations in WM provides a strong complement to the extant work.

The term mental imagery has been used to refer to two different cognitive operations. One is acting on representations in a way to transform them, such as mental rotation (Albers et al., 2013; Christophel et al., 2015; D’Esposito et al., 1999; Glahn et al., 2002). This type of mental imagery is the focus of the present manuscript. The term mental imagery has also been used to refer to recalling and imagining previously learned stimuli (Buchsbaum et al., 2012; Naselaris et al., 2015; Polyn et al., 2005; Slotnick et al., 2005; Xie et al., 2020). Overall, these studies showed that mental imagery, whether transformation or retrieval based, is established via neural activation of early perceptual visual regions as WM storage does (but see Knauff et al., 2000). Moreover, recent MVPA studies showed that not only the level of activity but also patterns of activity in these regions overlapped in imagery and perception (Albers et al., 2013; Christophel et al., 2015). However, all but one of these studies investigated mental imagery in the visual domain, using oriented lines, faces, objects, or abstract shapes. The only study that investigated the transformation of spatial information used univariate methods (Glahn et al., 2002). Although this study showed that there was an overlap in the brain regions that were active during storage and transformation in spatial, it was not able to show whether the information content was represented in overlapping brain regions. Even so, representations in the same brain areas do not imply overlapping representations (van Loon et al., 2018; Yu et al., 2020). Thus, our results provide the first evidence of overlapping neural representations of WM storage and mental imagery in the spatial domain.

Previous studies found enhanced activity in the dorsolateral prefrontal cortex for transformation vs. storage in WM using fMRI (D’Esposito, Postle, Ballard, & Lease, 1999; Glahn et al., 2002; Veltman, Rombouts, & Dolan, 2003). These results suggest that transformation might require additional attentional resources compared to mere storage in WM. This raises the possibility that transformed and static WM representations might be indexed by activity in different frequency bands, given past work finding shifts in the frequency bands containing information about prioritized and unprioritized memory representations (Rose et al., 2016). Nevertheless, we found that the same spatially-selective patterns of alpha activity tracked the initial and transformed positions, despite the requirement to actively manipulate the stored information.

The CTF slope for the transformed position during the 2^nd^ retention interval predicted RT across subjects (Experiment 1), with increased selectivity predicting faster manual report of the transformed position. Given that subjects were not instructed to respond as quickly as possible in this experiment, our speculation is that this finding may reflect strategic differences between subjects regarding the timing of their mental rotation. Some subjects may have adopted a more prospective strategy of immediately shifting the representation to the cued position, while other subjects may have waited until closer to the end of the delay period to perform the mental rotation. Thus, our hypothesis is that subjects who rotated early had more time to prepare their response before the test display was presented. Indeed, our switch to a speeded task in Experiment 2 was motivated by the correlation observed in Experiment 1. When subjects in Experiment 2 were explicitly instructed to respond as fast as possible, the average RT was ∼3 times faster, and the standard deviation was 4.9 times smaller compared to Experiment 1. This difference in RT between the two experiments that differed in instructions suggests that the rate with which WM transformation takes place can be adjusted based on incentives and strategies. This conclusion is consistent with a recent finding that showed the use of a prospective strategy in WM could be biased by task demands (Lewis-Peacock et al., 2016). Having a large number of trials in Experiment 2 (almost double of Experiment 1) allowed an analysis across trials using a median split approach on RT. The location specificity for the transformed content, as indexed by the CTF slope, was higher in fast RT trials than in slow RT trials. This result is in line with the between-subject correlation between average RT and average CTF slope in Experiment 1 and strengthens our conclusion that alpha activity tracks the temporal dynamics of this mental rotation task.

To conclude, we show that transformed and static representations in spatial WM are represented in overlapping neural networks as reflected in the topography of alpha-band (8-12 Hz) power recorded via EEG. These findings highlight a useful approach for tracking endogenous transformations of online memory representations.

## Acknowledgments

This work was supported by NIMH grant 2R01MH087214-06A1. We would like to thank Shannon Heald for creating the sound files used as the transformation cues, and Ariana Gale, Brendan Colson, Melanie Sykes, and Emily Xue for assisting with data collection.

## Supplementary Material

**Supp Fig 1.**
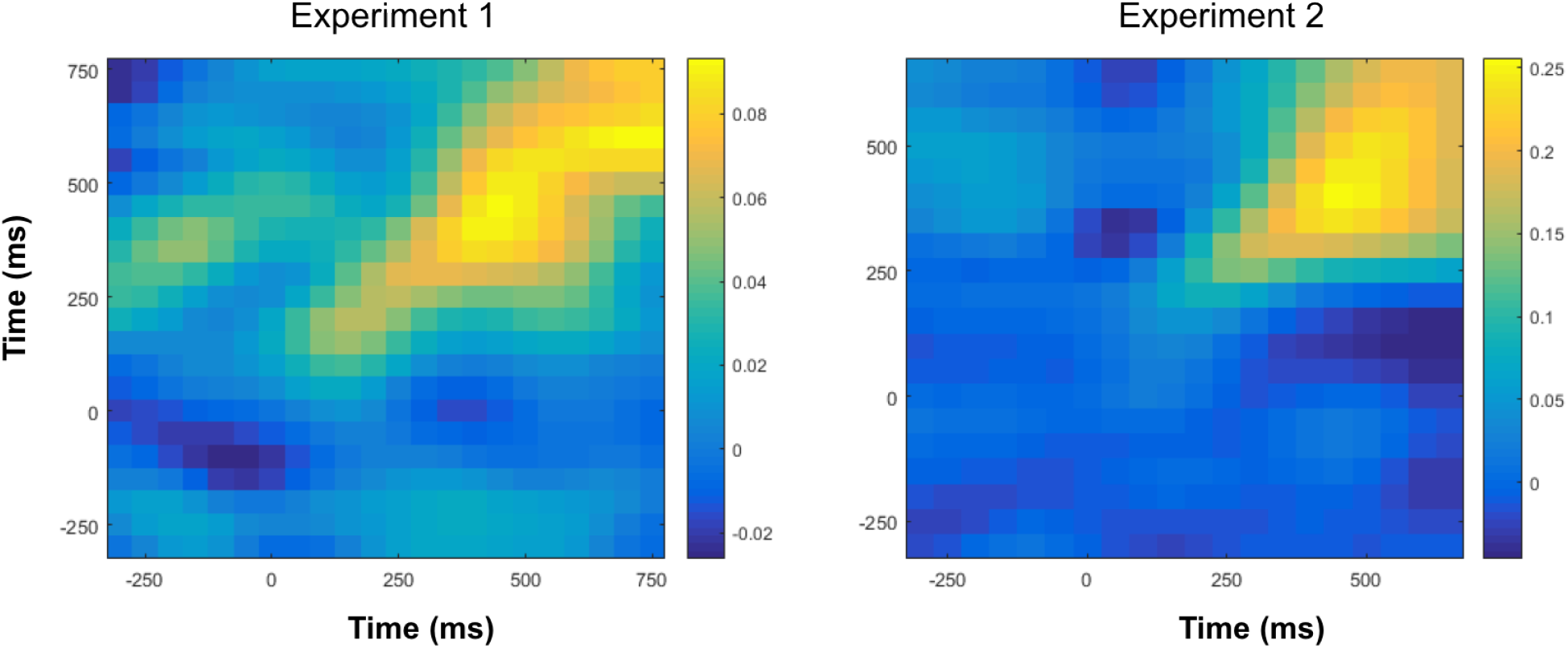
Cross-temporal CTF slope prior to the onset of the transformation cue, separately for Experiment 1 (left) and Experiment 2 (right). The CTF slope becomes stable after about 300 ms in both experiments.

**Supp Fig 2.**
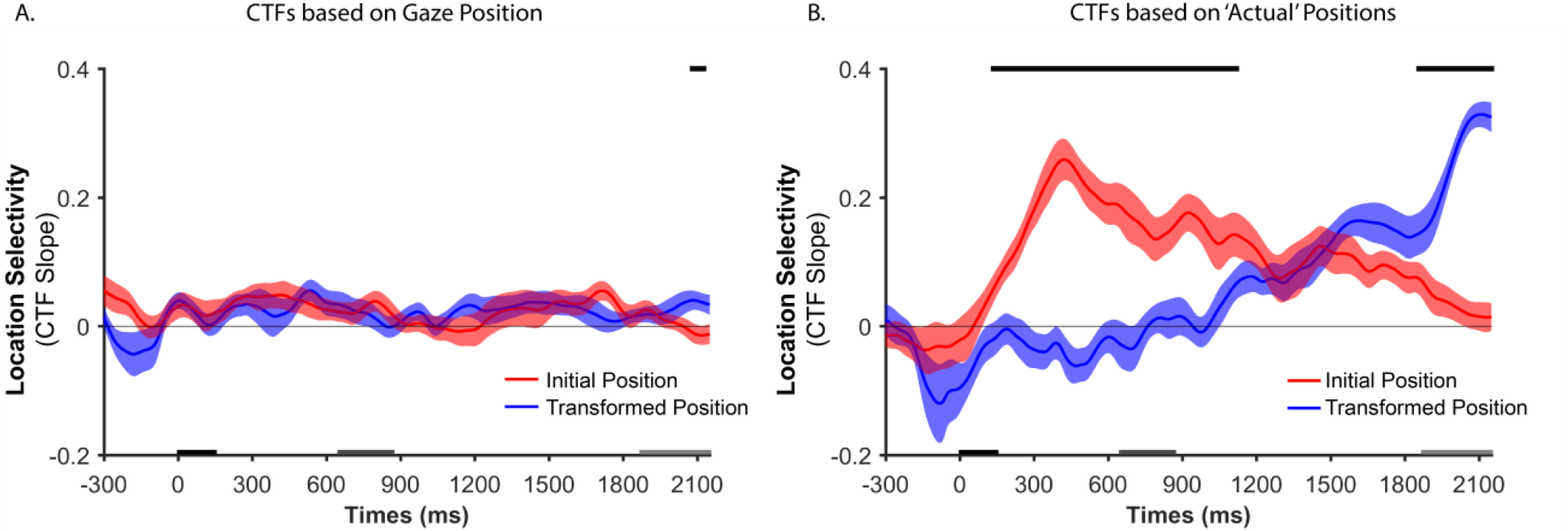
Location selectivity in the EEG that is quantified as CTF slopes for Experiment 2 that are calculated using position bins based on **(a)** gaze position and **(b)** actual WM positions as transformed in the experiment. Initial and transformed positions are shown in different colors. The black markers along the top of the panes indicate the time points at which the CTF slopes were reliably different between initial and transformed positions as indicated by a cluster-based permutation test (p*<0*.*05;* two-tailed). Shaded error bars show the bootstrapped standard error of the mean CTF slope. These results show that eye movements cannot account for the location selectivity in the EEG signal.

